# A *substrate* for modular, extensible data-visualization

**DOI:** 10.1101/217349

**Authors:** Jordan K Matelsky, Joseph Downs, Hannah Cowley, Brock Wester, William Gray-Roncal

## Abstract

As the scope of scientific questions increase and datasets grow larger, the visualization of relevant information correspondingly becomes more difficult and complex. Sharing visualizations amongst collaborators and with the public can be especially onerous, as it is challenging to reconcile software dependencies, data formats, and specific user needs in an easily accessible package. We present `substrate`, a data-visualization framework designed to simplify communication and code reuse across diverse research teams. Our platform provides a simple, powerful, browser-based interface for scientists to rapidly build effective three-dimensional scenes and visualizations. We aim to reduce the gap of existing systems, which commonly prescribe a limited set of high-level components, that are rarely optimized for arbitrarily large data visualization or for custom data types. To further engage the broader scientific community and enable seamless integration with existing scientific workflows, we also present `pytri`, a Python library that bridges the use of `substrate` with the ubiquitous scientific computing platform, *Jupyter*. Our intention is to reduce the activation energy required to transition between exploratory data analysis, data visualization, and publication-quality interactive scenes.

## Introduction

Using modern web-based visualization frameworks [1, 2] makes it easy to generate beautiful, interactive, and informative visualizations of scientific data. These renderings simplify the processes of exploring data and sharing insights with the community. In many domains, this has become a key step in the research and discovery pipeline [3].

One challenge with these technologies is the difficulty of adapting prior visualization work to a new use case. These tools are often built to be single-purpose rather than interoperable. Therefore it can be difficult or even impossible to combine aspects of disparate visualization scenes, even when the visualizations use the same technologies or frameworks. This challenge leads to software duplication instead of reuse, and complicates the portability of these products between research efforts. Often, modern visualization systems have chosen to either enjoy wide adoption at the expense of domain-specific tooling (e.g., plotly and matplotlib), or have focused on scientific subdomains at the expense of extensibility (e.g., common GIS or biological rendering software such as neuroglancer [4] and FreeSurfer [5]). As a result, combining visuals from more than one analysis or modality often requires significant engineering effort [6].

Scientists and software developers have produced several frameworks designed to remedy these challenges [[3, 7, 8] and we leverage architectural ideas from some of these frameworks in our solution, called `substrate`. We follow a compositional model similar in spirit to component-based engines such as React or Vue [[8, 9, 4, 10], wherein a visualization is comprised of many individual, independent components. Furthermore, we have developed `pytri`, a Python module that enables Python developers to access and interact with WebGL-based `substrate` from inside a Jupyter environment without requiring prior JavaScript or WebGL knowledge. This ecosystem includes integrated Jupyter notebook capabilities found in systems such as Mayavi or Neuroglancer [7, 4] alongside generalized visualization capabilities found in systems such as Plotly or Matplotlib [11, 12].

Unlike many existing Jupyter visualization packages, `pytri` visualizations are fully customizable, even by the end user. Users are not constrained by the limits of prepackaged visualization data structures or plot types, and can combine prebuilt components alongside custom, purpose-built visualization components. Users directly interact with their underlying data as usual, while these tools bring visualization capabilities fully into the data analytics platform.

Our tools are designed to enable the visualization of large-scale data or custom datatypes, coregister multimodal data, and simplify the process of sharing or reproducing analyses — all without disrupting the data science process.

We have designed our approach to be usable with minimal technical knowledge: A user can enter this ecosystem with only a basic knowledge of JavaScript or Python, though power-users and proficient software developers will find these libraries easy to manipulate and extend to suit their needs.

We first describe the software design of `substrate`, and then introduce `pytri` in order to render `substrate` scenes from common Python data libraries such as numpy [13], networkx [14], or pandas [15]. Finally, we share example use-cases in which the interoperability provided by `substrate` can reduce the engineering overhead of a new visualization project. Code and tutorials are available online at the links in the *Availability of Data and Material* section.

## 1 Architecture

To fully separate the responsibilities of `substrate`, our scene engine, and `pytri`, our Python integration library, our software architecture mandates that all rendering and Document Object Model (DOM) manipulation is handled by `substrate` and all Python manipulations and datatype translation take place in `pytri`.

### substrate

`substrate` is a JavaScript library that exposes a simple but powerful developer-facing application program interface (API). This abstraction enables otherwise disparate visualization projects to share resources and logic easily. The `substrate` JavaScript library is intended to simplify component-reusability of commonly used data-visualization structures such as scatter plots or 3D-embedded graph visualizations. Each atomic visualization component implements a common `Layer` API. Each `Layer` in a scene handles its own data management and interactivity in isolation, while a parent `Visualizer` object manages a list of these `Layers` in much the same way as a React application re-renders its components as needed when its internal state changes [9]. For example, a `ScatterLayer` implements `Layer` by accepting an array of [*x, y, z*]-tuples, and it will render these data when initialized. In the same scene, a `GraphLayer` might render a 3D graph embedding, as shown in Figure 1. These layers exist in the same 3D coordinate system, but are managed independently so changes to one does not require a re-render of the other components in the scene. Comprehensive `Layer` API documentation is available online.

**Figure 1:**
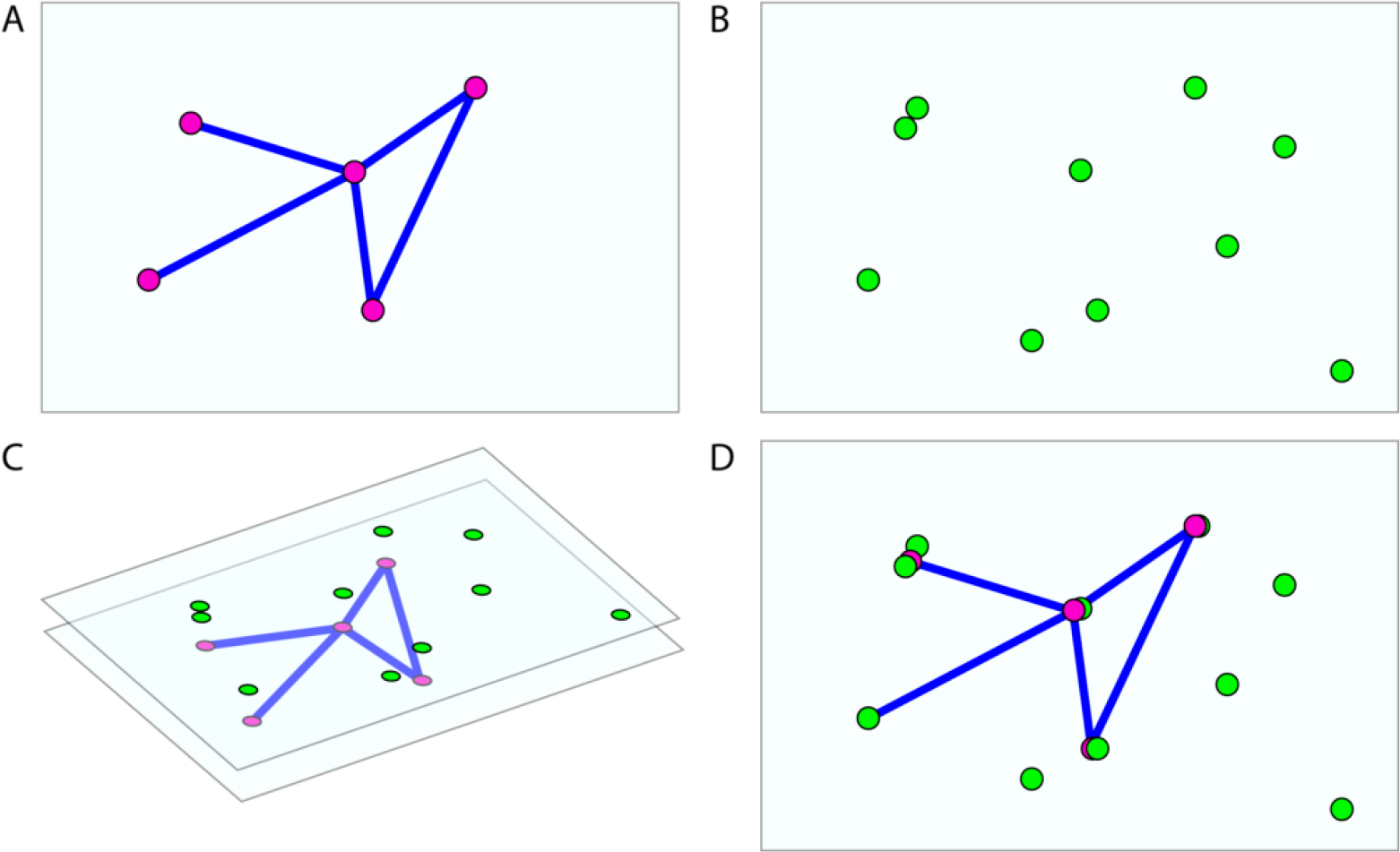
A `GraphLayer` and a `ScatterLayer`, embedded in the same 3D space. `substrate` overlays the two and controls the two independently. Ⓐ One layer renders a graph object. Ⓑ A second layer renders a scatter of points in 3D space. Ⓒ The layers are composited by the substrate Visualizer into a single image. Ⓓ The rendered scene, viewed from the substrate camera in finished form.

We formalize a universal `Layer` interface that accommodates common visualization tasks. `Layers` must include:

1. `requestInit` (function) is called before the visualization starts: This function generally includes instructions to provision objects in a 3D scene, or to request data from a remote source with a long round-trip
2. `requestRender` (function) runs on every frame. In static, non-animated `Layers`, this function may be empty or remain unimplemented in order to conserve compute power
3. `children` (array attribute) lists all objects in a scene associated with a particular `Layer`. When a `Layer` is removed from the visualization, all objects in this list are cleaned up by `substrate` internally.

This simple interface enables many different visualization objects or groups of objects to coexist in a scene without interfering with one another. Such namespacing conflicts are a common pitfall when combining conventional, separately-developed three.js assets into a single scene.

In order to improve the accessibility of our codebase to new developers and data scientists, we use *three.js* as a convenience to wrap WebGL. Despite the prevalence of *three.js* in our current codebase, `substrate` aims to be framework-agnostic. Authors of new `Layers` may choose to write WebGL directly, or use another wrapper or framework. `substrate` will support these `Layers` provided they subscribe to the `substrate.Layer` and `substrate.Visualizer` interfaces. In this way, `substrate` works in a similar extensible fashion to the Uber `deck.gl` project, though deck.gl reimplements full scene rendering from the ground up, and `substrate` uses the *three.js* industry standard [2].

The compositional property of `Layers` can be expressed in code using the syntax shown in Figure 2. Here we show a simple `Visualizer` containing two `Layers`; this short snippet can run a complete visualization without any extra configuration.

**Figure 2:**
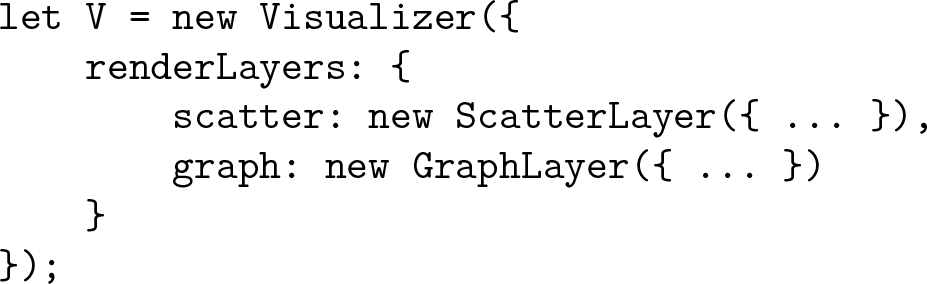
A sample `Visualizer`, with two `Layer`s. One renders a 3D scatter-plot, and the other renders nodes and edges of an undirected graph.

One common use of separate `Layers` is to place objects of interest — such as a mesh — in one `Layer`, and place lighting, axes, or other environmental factors in another. This enables a researcher to share their core data, such as the 3D mesh, with other researchers, without including extraneous features such as light sources or grids and axes. (This simultaneously enables an artist to reuse their lighting layout across projects.) In Figure 3, we illustrate a sample `Layer` implementation that can be ported to any `substrate` visualization. Our `add-and-remove-layer` demo provides an example in which a `MeshLayer` is added or removed, without affecting other objects in the scene.

**Figure 3:**
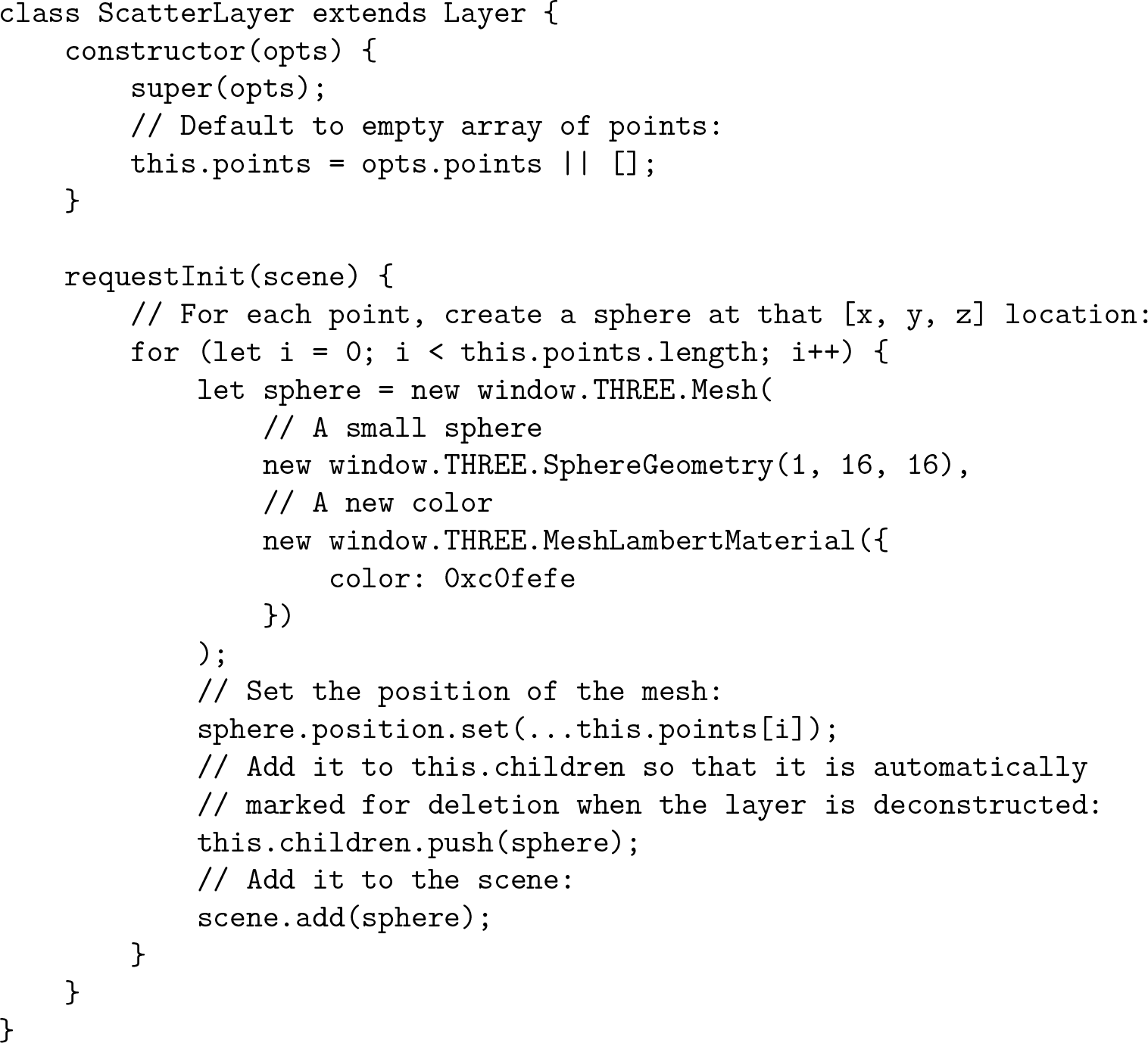
A sample implementation of a `Layer` that generates a point-cloud from the data provided in the constructor. This *exact implementation* can be dropped into any `substrate` visualization without modification. This code, and other `Layer` examples, are available online.

`Layers` written for one visualization or application may be repurposed or reused in another scene without additional developer effort. This means that for most visualization usecases, such as graph displays or scatter plots, no `substrate` knowledge is required at all; instead, prebuilt `Layers` are available for public use, including a `ScatterLayer`, `GraphLayer`, `MeshLayer`, and many others, as seen in Table 1.

**Table 1:** Available layers built into the substrate package.

| Layer name | Description | Useful for... |
| --- | --- | --- |
| AxisLayer | Contains an RGB axis at the scene origin | ...orienting the user in space |
| GraphLayer | A graph renderer with support for many millions of nodes or edges | ...complex networks |
| ImageLayer | Import 2D images and render them as a plane | ...displaying static iamgery |
| LightingLayer | Three-point lighting scheme, as commonly used in film and 3D graphics | ...easy lighting setups |
| LineSegmentsLayer | A list of line segments to be rendered individually | ...paths or wireframes |
| MeshLayer | Arbitrary 3D meshes, imported in common file formats such as OBJ | ...arbitrary 3D solids |
| ScatterLayer | Point-clouds of many millions of points | ...colored scatter plots |
| VolumeLayer | A visualization of dense 3D voxelwise data | ...rendering volumetric data |

In some cases, specific data requirements may mean that researchers cannot use these existing, prebuilt `Layer` implementations. If a developer decides to implement their own `Layer` from scratch, it can be trivially integrated into other visualizations, as all `substrate Layers` subscribe to the same simple interface and are interchangeable. For example, the bigdata neuroscience community has developed a custom layer that visualizes larger-than-RAM imagery by shuttling data in and out of memory as it enters and leaves the `substrate` camera viewport. Social graph research teams have developed representations of graphs with enriched visual cues to signal node and edge attributes.

When research usecases require customization, engineers can easily implement `Layers` that suit their specific need. Prebuilt `Layers` are sufficient for many applications, and require no engineering knowledge or ability from the end-user.

Our `brownian-particle-motion` example demonstrates how a developer can easily implement a custom `Layer`, while still taking advantage of prebuilt code. We envision that community users may request to merge commonly used `Layers` into the `substrate` codebase to extend native functionality and cover a diverse set of usecases. As groups work together to achieve research goals, these researchers may separately develop `Layers` (e.g. a raw experimental `Layer` and an annotation `Layer` for the analysis) which can be combined in the same scene when needed.

### pytri

Requiring a scientist to exit their research environment in order to engage with visualizations reduces the time-efficacy of research [16, 17]. In order to provide a convenient, inline visualization solution for data scientists, we created `pytri`, a Python package that enables visualization of `substrate Layers` without leaving a Jupyter notebook [18] or other IPython environment (Figure 4). Jupyter is a standard research platform for many communities. By bringing composable, extensible visualization to this platform, data scientists can quickly visualize and explore data in a familiar environment without needing to understand the underlying `substrate` codebase.

**Figure 4:**
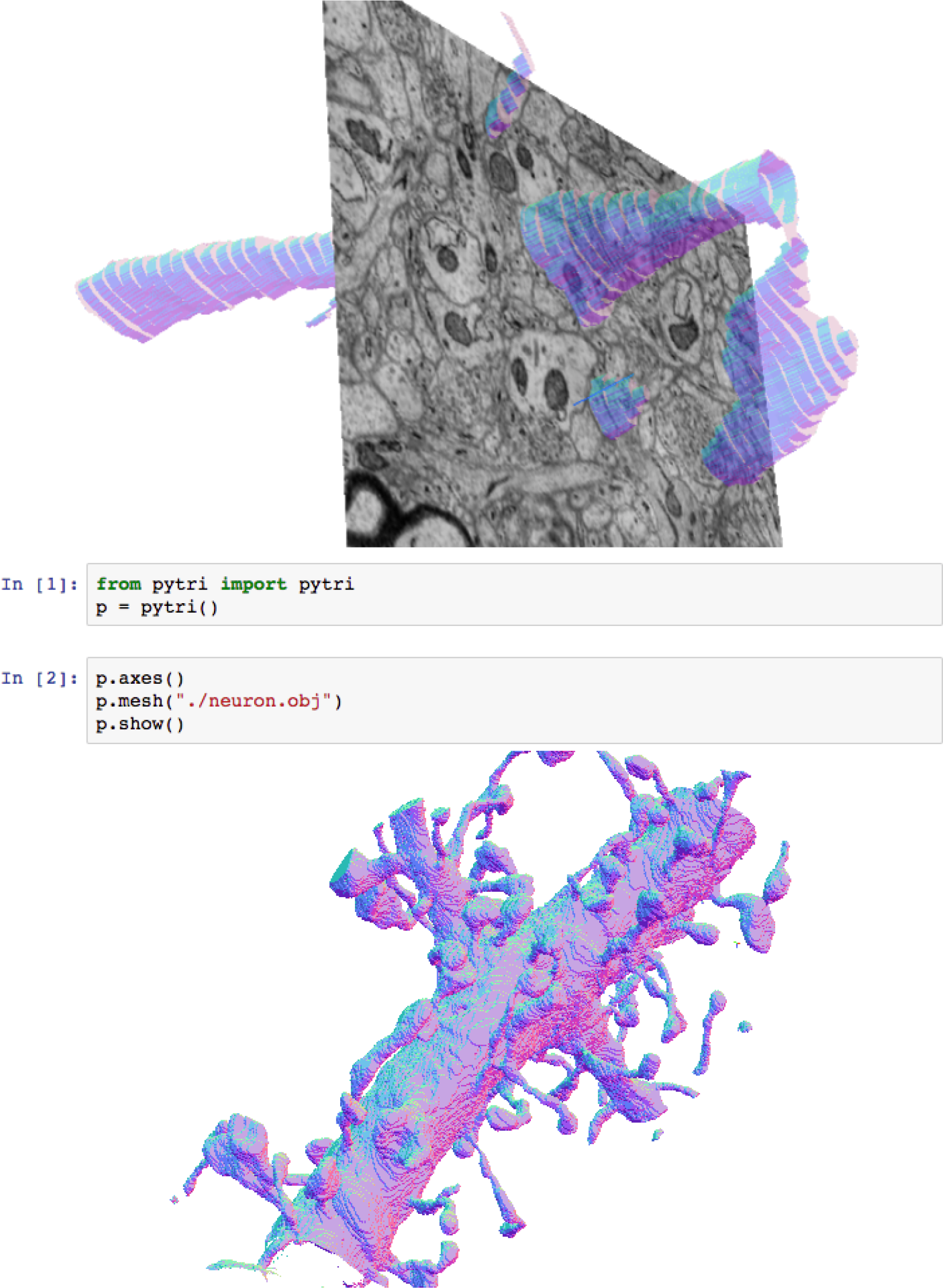
Here, `pytri` runs `substrate` visualizations inside a Jupyter notebook, using WebGL to enable real-time interaction with the visualization. This high-detail mesh was generated using manual annotations from a recent electron microscopy study [30].

**Figure 5:**
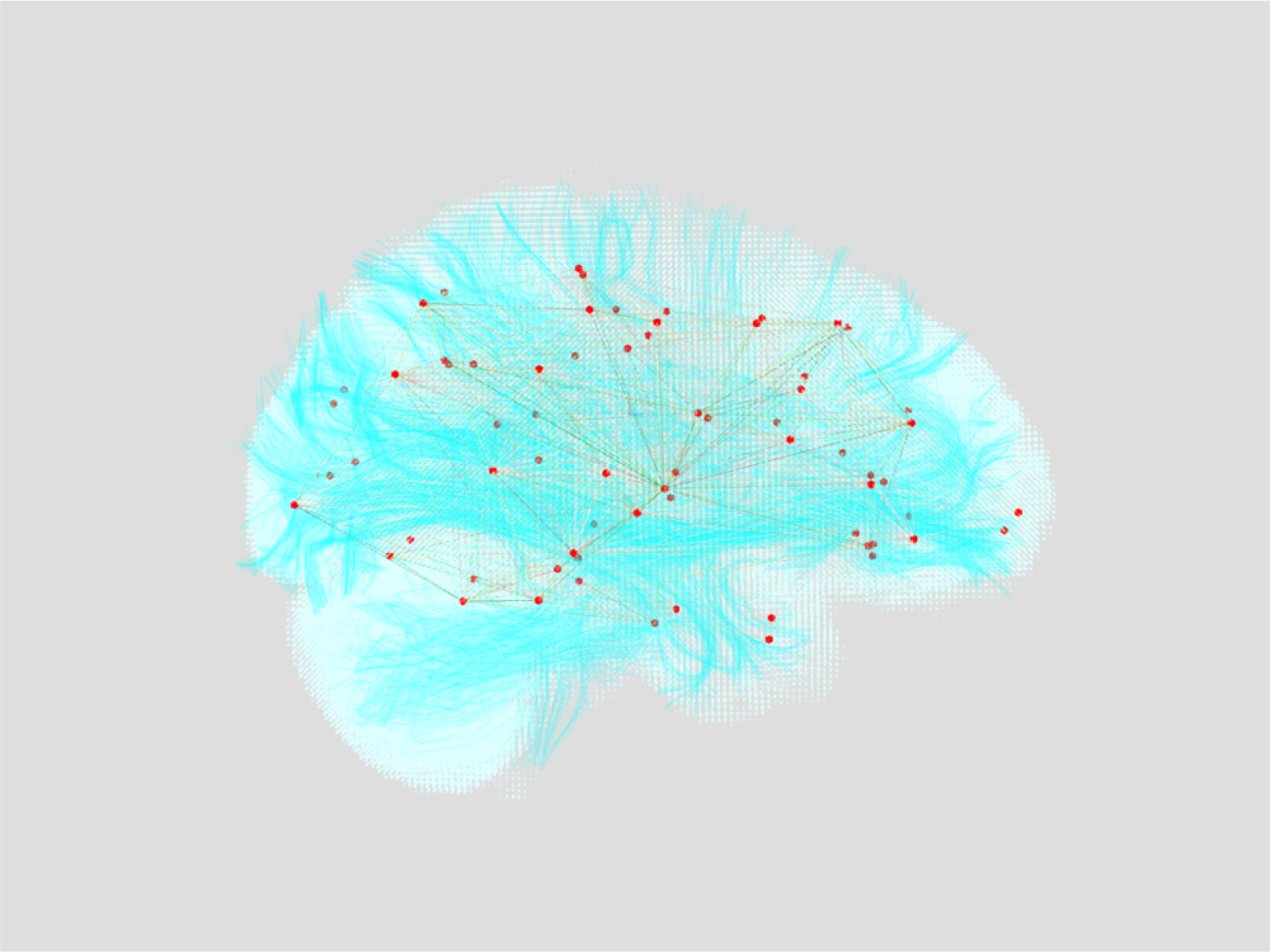
A view of the data using volumetric rendering. A graph of estimated brain region connectivity is rendered in red and yellow. Estimated fibers are rendered in cyan. The white brain volume is the direct output of DTI MRI.

**Figure 6:**
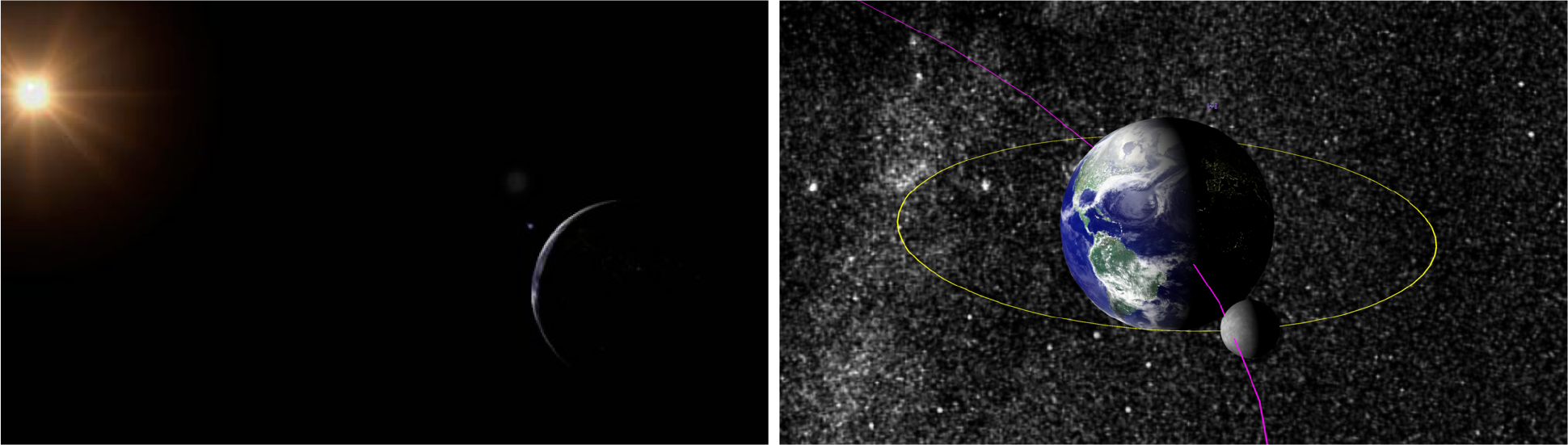
We use `substrate` to render the orbits of the Earth and Moon around the sun according to their realtime positions. A mesh representation of the International Space Station pulls live positional data from the internet [21, 22]. Orbital body sizes and trajectories are exaggerated for visibility.

Our initial use-case demanded performant largescale graph visualization. Though standalone visualization software existed for this scale of graph, our team found it onerous to exit our data science platform to view the raw data. Furthermore, we encountered issues with the performance of many popular 3D plotting libraries in Python, as they were unable to handle the size of graphs we needed to visualize. We built `substrate` to gracefully handle graph data containing many millions of edges. We then added corresponding hooks in `pytri` to empower users to create large-scale visualizations quickly without leaving the familiar Python environment.

`pytri` combines `substrate` capability with Python datatypes by leveraging the Jupyter Notebook `IPython.display` module, to which `substrate` delegates responsibility for interaction with the Jupyter DOM through both JavaScript (`IPython.display.JavaScript`) and HTML (`IPython.display.HTML`) [18].

These DOM manipulations, unlike the current Jupyter Widget architecture, enable `pytri` visualizations and data to persist even in a static HTML export of the Jupyter notebook. This enables the distribution of reproducible visualizations without requiring the end-user to install or configure software packages. In other words, researchers can produce static HTML files with interactive data visualizations which can be shared by email, by online publication, or through sharing the original source code.

## 2 Results

The following use cases illustrate `substrate`’s flexibility, idiomatic brevity, and ability to handle custom data. By lowering the overhead associated with task-switching between visualization and analysis, `substrate` provides an opportunity for a team to more intimately explore their data and iterate on analyses in realtime.

One of the key advantages of `substrate` is its use as a general framework for visualization across many domains. Here we highlight a few diverse applications that benefit from this work.

### 2.1 Use Case: Analyzing Biological Imaging Data

Biomedical research is one of many domains that benefits from the recent explosion in big data [19]. As such, there is pressing need for research tools that can adapt to the demands of large-scale datasets. Such a customizable and interactive framework for visualization can help researchers understand otherwise uninterpretable data. Volumetric imagery is one datatype that is particularly important both in biomedical research as well as in the clinical setting. Many existing frameworks for data visualization lack a lightweight tool that preserves the vital spatial relationships found in volumetric imagery.

When creating visualizations using `substrate` and `pytri`, the researcher has the speed and flexibility necessary to transform a typical visualization product from a static communication tool to a dynamic exploration aid. As an exercise in using `pytri` for data exploration and scientific communication, we present the following use cases, showcasing similar opportunities across six orders of magnitude (e.g., magnetic resonance imaging (MRI) to electron microscopy (EM)). We focus on an exciting area of contemporary research called connectomics, which is a domain focused on building brain connectivity maps at various resolutions.

We first investigate slices of MRI data volumes using `ImageLayer`. Navigation through the dense 3D data can be done programmatically through direct calls to the `pytri` API, or through the `Jupyter` UI, so the researcher never has to leave their data analysis environment. Beginning with this volumetric data representation, we can overlay fiber tracts representing estimates of major axonal bundles in the brain, and the derived nodes and edges associated with a connectome [20]. More specificially, the connectome is visualized using a `GraphLayer` overlaid on a `MeshLayer` of the surface of the patient’s brain as computed from structural MRI. When the analysis is ready to be shared, the researcher can easily package these visualizations for others by exporting the Jupyter notebook to an interactive HTML page.

For EM data, we show the flexibility of the tooling by displaying imagery slices, along with commonly used derived data representations such as meshes generated from manual or automated segmentation methods and skeleton traces. These tools are important to rapidly explore and validate large reconstructions. Because our visualization tools exist within an analytics environment, users can compute quantitative analyses in the same environment, reducing impediments to discovery. `substrate`’s layer-based framework allows the researcher to overlay multiple data sources, to reorient the view to focus on one detailed section of the data at a time, and to fully leverage data visualization in the research and sharing process.

### 2.2 Use Case: Astronomical Observational Data

The civilian space community communicates mission details to a diverse audience, and visualization greatly enables public and collaborator understanding of mission planning and execution. The ability to quickly produce a visual summary of the mission can also enable plan iteration and facilitate a discussion of alternative solutions. 3D visualization of a mission can assist with exploring the complex maneuvers sometimes demanded in space exploration.

`substrate` is well-suited to display orbital and hyperbolic trajectories of bodies moving through outer space. In 6, we show the paths and bodies of the Earth/Moon system and the International Space Station (ISS) using only a `FiberLayer` to represent orbital paths and a `MeshLayer` (to render a downloaded 3D mesh of the ISS [21]) in `substrate`. The ISS position is updated in realtime to its current real-world position using the `requestRender` function of the `Layer` API (with data pulled from an online resource [22]). The sizes of the orbital bodies or satellites can be changed easily by removing, resizing, or reinserting the corresponding meshes. Many possible trajectories can be viewed in rapid succession by toggling their visibility in the scene. This usecase provides an example of how existing tools might be augmented through a simple, web-based visualization environment to engage the public and produce publication-ready graphics for community consumption.

### 2.3 Use Case 3: Geospatial Information Systems

Geospatial information is of interest to researchers in a variety of domains, including agriculture, architecture, and urban planning. Many of the existing state of the art geographic information systems (GIS) require standalone software installations, and visualizations are often handled in a separate application than that in which the initial data science is performed [23]. This requires researchers to switch between analysis and exploration, or else it constrains research pipelines to live inside of specialized visualization software such as QGIS [24] or SAGA [25].

Using `pytri`, GIS data can be visualized natively in Jupyter alongside analyses, and users may then visually explore the byproducts of this exploration without leaving the Jupyter notebook. Here, we perform a basic query of GIS data in the Johns Hopkins University Homewood Campus area using the `osmnx` Python package [23], one example of a tool one might use to download a large-scale graph. We then demonstrate the ability to coregister the visualizations of a graph of street connectivity alongside regions of interest and structure meshes downloaded from *3D Warehouse* [26, 27]. This provides a flexible framework to enrich a scene as additional sensors and data fusion products become available. We use `pytri` to visualizate these data science products in a Jupyter notebook in Figure 7.

**Figure 7:**
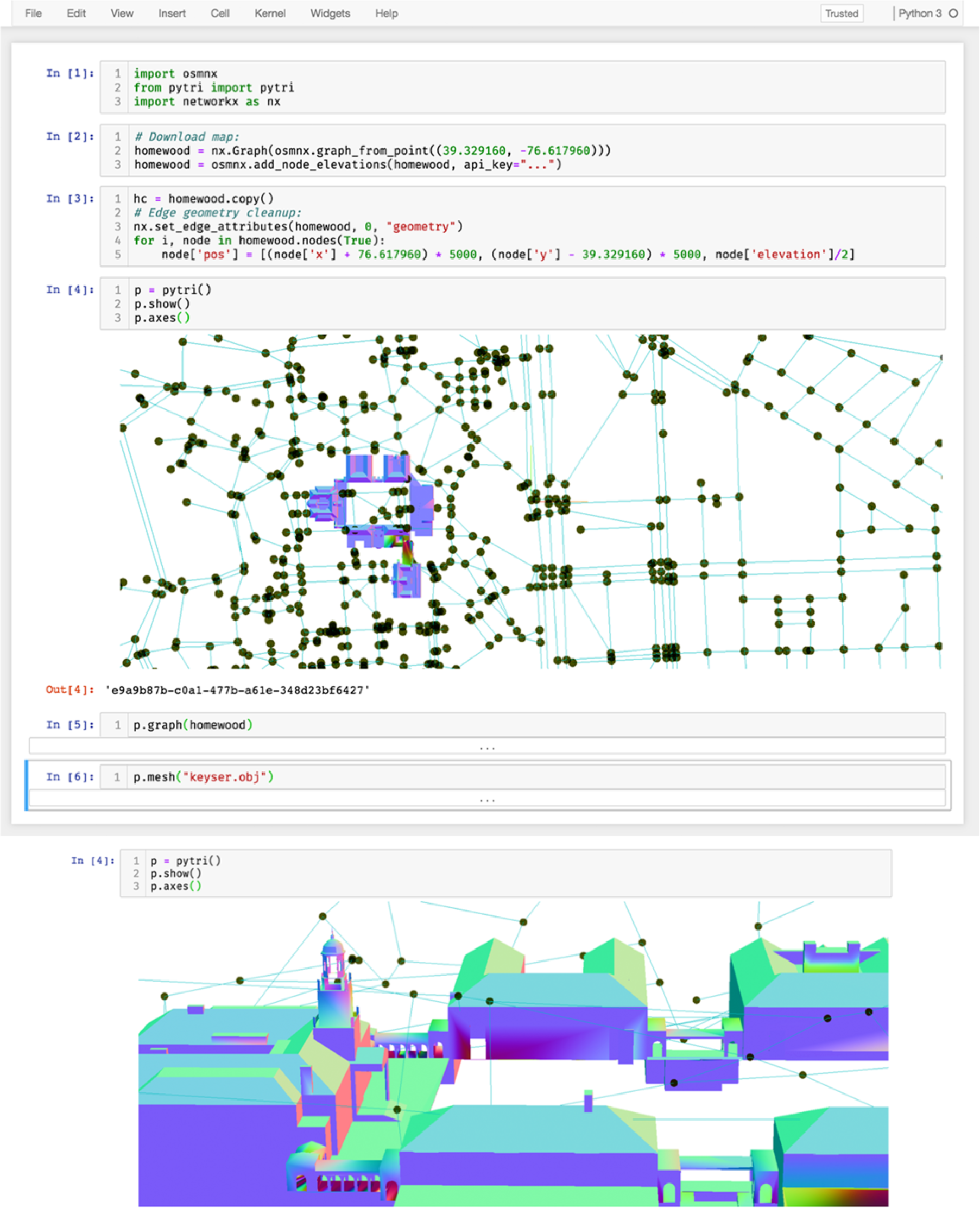
Using `osmnx`[23], `pytri` displays a graph representation of the roads and paths surrounding the Johns Hopkins University Homewood campus. **Top**: The researcher never leaves the Jupyter notebook. **Bottom**: Clicking and dragging the visualization yields a different view of the Gilman Clock Tower. No re-render is required: The user can interact with the visualization in realtime.

This visualization uses a `GraphLayer` to represent streets, paths, and intersections as generated by the osmnx library in `networkx.Graph` format [14]. A `MeshLayer` is used to overlay a rendering of the Keyser Quad buildings in the same 3D coordinate frame.

Existing GIS visualization software tools often require that analysis is performed offline and data products are ingested — or else analyses are constrained to software-specific plugin architectures. The use of `pytri` enables simple 3D coregistered visualization of multimodal geographic data in a variety of data formats, including networkx graph and OBJ-formatted meshes.

## 3 Discussion

Similar to how web frameworks such as Angular [28], React [9], and Vue [10] popularized the *reusable component* model for maintainable interface composition, our work emphasizes the reusability of *visualization* components by exposing an interface for discrete entities in a 3D scene. By combining several of these components, complex and deeply informative scenes can be designed with only minimal engineering effort. We share our implementation of this common interface in our software package `substrate`, which we provide as an open-source resource to the scientific community, and in `pytri`, a Python software package that enables the use of this common API without advanced JavaScript knowledge. We intend that researchers unfamiliar with visualization technology will use `pytri` to compose visualizations in-line with their analyses, thus removing some of the activation energy required to produce and share reproducible, publication-ready graphics.

The list of components we provide is nonexhaustive, and intend to support the community in efforts to increase the breadth of domains aided by `substrate`. Because many users may choose to conduct data science or research in languages besides Python, we intend to develop libraries for other common data science languages such as R and Julia, based upon ongoing community feedback.

Efforts are ongoing to natively support very large (out-of-RAM) dataset representations in both `substrate` and `pytri` libraries. Users with different tooling requirements, who require custom import formats, or who rely upon very large scale visualizations (e.g. graphs with billions of vertices and edges) may require additional engineering effort to fully leverage the `substrate` ecosystem.

As scientific datasets grow in size and complexity, communicating relevant data clearly and effectively is more important now than ever. Large-scale, multiteam efforts require portable and shareable visualizations that can be developed by several engineers simultaneously, and used by entire teams of both technical as well as non-technical individuals. It is our hope that tools like `substrate` and `pytri` will help support reproducible, reusable scientific discovery in the data science community. Our code and data are publicly available as described in the *Availability of Data and Material* section.

## Availability of Data and Material

We provide the codebase for substrate, documented and open-source under the Apache 2.0 License at https://github.com/aplbrain/substrate, and welcome community feedback in the form of pull requests, feature suggestions, or bug reports. We also provide demonstrations of common uses and tutorials for users to extend the current functionality. Finally, we provide a Dockerfile to allow anyone to trivially launch a `pytri`-enabled Jupyter notebook in their browser [29]. `pytri` can be downloaded either via `pypi` (`pip install pytri`) or from our open-source repository at https://github.com/aplbrain/pytri. Demonstrations of use-cases, as well as explanatory code included in this manuscript, are available at our separate repository, https://github.com/aplbrain/substrate-demos. Comprehensive substrate documentation is available online at https://aplbrain.github.io/substrate/.

